# Genetic tools weed out misconceptions of strain reliability in *Cannabis sativa*: Implications for a budding industry

**DOI:** 10.1101/332320

**Authors:** Anna L. Schwabe, Mitchell E. McGlaughlin

## Abstract

*Cannabis sativa* is listed as a Schedule I substance by the United States Drug Enforcement Agency and has been federally illegal in the United States since 1937. However, the majority of states in the United States, as well as several countries, now have various levels of legal *Cannabis*. Products are labeled with identifying strain names but there is no official mechanism to register *Cannabis* strains, therefore the potential exists for incorrect identification or labeling. This study uses genetic analyses to investigate strain reliability from the consumer point of view. Ten microsatellite regions were used to examine samples from strains obtained from dispensaries in three states. Samples were examined for genetic similarity within strains, and also a possible genetic distinction between Sativa, Indica, or Hybrid types. The analyses revealed genetic inconsistencies within strains. Additionally, although there was strong statistical support dividing the samples into two genetic groups, the groups did not correspond to commonly reported Sativa/Hybrid/Indica types. Genetic differences have the potential to lead to phenotypic differences and unexpected effects, which could be surprising for the recreational user, but have more serious implications for patients relying on strains that alleviate specific medical symptoms.

## Introduction

*Cannabis sativa* L. is one of the most useful plants (Clarke & Merlin, 2013) with evidence of human cultivation dating back thousands of years (Abel, 2013). *Cannabis* prohibition in the United States began with the Marihuana Tax Act in 1937 (The Marihuana Tax Act of 1937), and the Controlled Substances Act of 1970 classified *Cannabis* as a Schedule I drug with no “accepted medical use in treatment in the United States” (Controlled Substances Act, 1970). *Cannabis* is largely illegal worldwide, but laws allowing *Cannabis* for use as hemp, medicine, and some adult recreational use are emerging (ProCon, 2016a). *Cannabis* is a multi-billion dollar crop, but global restrictions have limited *Cannabis* related research. The origins and genetic identities of many *Cannabis* strains are largely unknown, as there are relatively few genetic studies focused on strains (Lynch *et al*., 2016).

The World Drug Report estimates ∼4.5% of the global population, consumes *Cannabis* regularly (United Nations Office on Drugs, Crime, 2010), and there are an estimated ∼3.5 million medical marijuana patients in the US (Marijuana Policy Project, 2017). Recent legalization has led to a surge of new strains as breeders are producing new plant varieties with novel chemical profiles with various psychotropic effects, and relief for an array of symptoms associated with medical conditions including (but not limited to): chronic pain, depression, anxiety, PTSD, autism, fibromyalgia, epilepsy, Chron’s Disease, and glaucoma (Ogborne *et al*., 2000; Tomida *et al*., 2004; Borgelt *et al.,* 2013; Naftali *et al.,* 2013; ProCon, 2016b).

Research using a variety of techniques consistently finds drug-types and hemp are genetically distinct (de Meijer *et al.,* 1996; Small, 1997; Sawler *et al*., 2015; Lynch *et al.,* 2016; Dufresnes *et al.,* 2017). Variation within the drug-types is higher than within hemp (Small, 1997; Sawler *et al.,* 2015; Lynch *et al*., 2016; Vergara *et al*., 2016). There is limited genetic research on variation within strains, but in studies with multiple accessions of a particular strain, variation is observed (Sawler *et al.,* 2015; Lynch *et al,*. 2016; Soler *et al.,* 2017).

There are generally two *Cannabis* usage groups (hemp and drug-types) although the scientific and common nomenclature is conflicted. The current Flora of North America recognizes all forms of *Cannabis* as *Cannabis sativa* L. (Small, 1997), but many breeders and botanists support the polytypic taxonomy of *Cannabis* based on morphological (de Lamarck & Poiret, 1789; Schultes, 1970; Emboden, 1974; Anderson, 1980), chemical (de Meijer *et al.,* 2003; Hillig & Mahlberg, 2004; Hillig, 2005; Hazekamp & Fischedick, 2012) and psychotropic (de Meijer *et al.,* 2003; Hillig & Mahlberg, 2004; Hazekamp & Fischedick, 2012; Clarke & Merlin, 2013) differences. However, the suggested putative species are presumed to readily interbreed and therefore violate species concepts that are applicable to plants (De Queiroz, 2007). The common terminology for *Cannabis* products are, that (1) hemp types have < 0.3% Δ9-tetrahydrocannabinol (THC), (2) plants of broad and narrow leaf drug-types as well as hybrid variants with moderate to high THC concentrations are referred to as marijuana, (3) drug-type strains of *Cannabis* are commonly divided into three categories: Sativa, Indica and Hybrid type strains, (4) drug-type strains with low THC and high cannabidiol (CBD) are sought after for medicinal use, and (5) there are thousands of variants of *Cannabis* referred to as strains. Genetic analyses have not provide a clear consensus for higher taxonomic distinction among these commonly described *Cannabis* types (Sawler *et al.,* 2015; Lynch *et al.,* 2016), but both the recreational and medical *Cannabis* communities claim there are distinct differences in effects between Sativa and Indica type strains (Smith, 2012; Leaf Science, 2014). Sativa type strains are associated with tall, loosely branched plants with long, narrow leaflets, and are reported to have energizing or uplifting psychotropic effects (Russo, 2007; Fischedick *et al.,* 2010; Hillig, 2004). Indica type strains are associated with shorter plants with dense branching and broad leaflets, and reportedly exhibit sedating effects and pain relieving properties (Russo, 2007; Fischedick *et al.,* 2010; Hillig, 2004). Hybrid types are a mix of varying degrees of the reported effects of Sativa and Indica types.

Morphological variation is typically used to categorize species, sub-species, and varieties. However, morphological identification can be difficult with closely related taxa and hybrid organisms (Rieseberg, 1995; Rieseberg, 1997; Cattell & Karl, 2004; Mallet, 2005; Zha *et al.,* 2008, Schwabe *et al*. 2015). Sexual reproduction generally results in offspring with a blend of traits from both parents. On the other hand, clonal offspring or progeny produced from self-fertilization should be virtually identical to the parent. Unique physical differences (phenotypes) and varying chemical profiles (chemotypes) may result when plants with the same genetic profile (genotype) are impacted by environmental factors (phenotypic plasticity) (Schlichting, 1986; Elzinga *et al*. 2015). Phenotypic plasticity is commonly observed in *Cannabis*, and therefore, the use of chemical profile or other physical characteristics are not ideal to precisely identify *Cannabis* variants (Schultes, 1970; Clarke & Merlin, 2013; Small, 2017)

Female flowers of predominantly dioecious *Cannabis* plants produce the majority of cannabinoids and terpenes in glandular trichomes. Female plants are selected based on desirable characters (mother plants) and are reproduced through cloning and, in some cases, self-fertilization to produce seeds (Green, 2005). The offspring will be identical (from clone), or nearly identical (from seed), to the mother plant. Cross-pollination allows for genetic variability and novel strain creation, but generally *Cannabis* growers use cloning to produce consistent products of established and popular strains. Whether propagated through cloning or from germination of self-fertilized seed, genetic variation within strains should be minimal no matter the source of origin.

There are an overwhelming number of *Cannabis* strains that vary widely in appearance, taste, smell and psychotropic effects (de Lamarck & Poiret, 1789; Schultes, 1970; Emboden, 1974; Anderson, 1980; de Meijer *et al.,* 2003; Hillig & Mahlberg, 2004; Hillig, 2005; Hazekamp & Fischedick, 2012; Clarke & Merlin, 2013). Strains are generally categorized as Indica, Sativa or Hybrid types. Online databases such as Leafly (Leafly, 2018) and Wikileaf (Wikileaf, 2018) provide consumers with information about strains but lack scientific merit for the *Cannabis* industry to regulate the consistency of strains. To our knowledge, there have not been any published scientific studies specifically investigating the genetic consistency of strains at multiple points of sale for *Cannabis* consumers.

Of particular interest is how the genetic integrity of named *Cannabis* strains over time in the absence of regulation been maintained (Green, 2014; Stockton, 2015). Other crop varieties are protected by certification through the United States Department of Agriculture (USDA) and The Plant Variety Protection Act of 1970 (PVPA), or similar mechanisms in other countries. This system protects against commercial exploitation, allows for trademarking, and recognizes intellectual property for developers of new plant cultivars (United States Department of Agriculture, 1989). Traditionally, morphological characters were used to define new varieties in crops such as grapes (*Vitis vinifera* L.), olives (*Olea europea* L.) and apples (*Malus domestica* Borkh.). With the rapid development of new varieties in these types of crops, morphological characters have become increasingly difficult to distinguish. Currently, quantitative and/or molecular characters are often used to demonstrate uniqueness among varieties to obtain a plant variety protection certificate from the Plant Variety Protection Office (PVPO) of the Agricultural Marketing Service, USDA (United States Department of Agriculture, 2015). Microsatellite genotyping enables growers and breeders of new cultivars to demonstrate uniqueness through variable genetic profiles (Rongwen *et al.,* 1995). Microsatellite genotyping has been used to distinguish cultivars and hybrid varieties of multiple crop varietals within species (Guilford *et al.,* 1997; Hokanson *et al.,* 1998; Cipriani *et al.,* 2002; Belaj *et al.,* 2004; Sarri *et al.,* 2006; Baldoni *et al.,* 2009; Štajner *et al.,* 2011; Costantini *et al.,* 2015; Pellerone *et al.,* 2015). Multiple crop studies have found that 3-12 microsatellite loci are sufficient to accurately identify varietals and detect misidentified individuals (Cipriani *et al.,* 2002; Belaj *et al.,* 2004; Sarri *et al.,* 2006; Poljuha *et al.,* 2008; Baldoni *et al.,* 2009; Muzzalupo *et al.,* 2009;). *Cannabis* varieties however, are not afforded any legal protections, as the USDA considers it an “ineligible commodity” (United States Department of Agriculture, 2016), but this system provides a model by which *Cannabis* strains could also be developed, identified, registered, and protected.

Currently, the *Cannabis* industry has no way to verify strains. Consequently, suppliers are unable to provide confirmation of strains. Reports of inconsistencies, along with the history of underground trading and growing in the absence of a verification system, reinforce the likelihood that strain names may be unreliable identifiers for *Cannabis* products at the present time. Without verification systems in place, there is the potential for misidentification and mislabeling of plants, creating names for plants of unknown origin, and even re-naming or re-labeling plants with prominent names for better sale. *Cannabis* taxonomy is complex, but given the success of microsatellites to determine varieties in other crops, we suggest the using genetic based approaches to provide identification information for strains in the medical and recreational marketplace.

Variable microsatellite markers were developed using the *Cannabis sativa* ‘Purple Kush’ draft genome (National Center for Biotechnology Information, accession AGQN00000000.1). These regions were compared within commercially available *C. sativa* strains to determine if products with the same name purchased from different sources have the genetic congruence we expect from propagation of clones or self-fertilized seeds. The unique approach for this study was that of the common retail consumer. Flower samples were purchased legally from dispensaries based on what was available at the time of purchase. All products were purchased as-is, with no additional information provided by the facility, other than the identifying label (strain name). This study aimed to determine if: (1) any genetic distinction separates the common perception of Sativa, Indica and Hybrid types; (2) purported proportions for Sativa, Indica and Hybrid type strains are reflected in the genotypes of multiple strains; (3) consistent genetic identity is found within a variety of different strain accessions obtained from different facilities; (4) there is evidence of misidentification or mislabeling.

## Materials and Methods

### Genetic Material

*Cannabis* samples for 30 strains were acquired from 20 dispensaries or donors in three states: Colorado - Denver (4), Boulder (3), Fort Collins (3), Garden City (4), Greeley (1), Longmont (1); California - San Luis Obispo (4); and Washington - Union Gap (1) (Table 1). All samples used in this study were obtained legally from either retail (Colorado and Washington), medical (California) dispensaries, or as a donation from legally obtained samples (Greeley 1). DNA was extracted using a modified CTAB extraction protocol (Doyle 1987) with 0.035-0.100 grams of dried flower tissue per extraction Proportions of Sativa and Indica phenotypes for each strain were retrieved from Wikileaf (Wikileaf, 2018). Analyses were performed on the full 122-sample dataset (Table 1). A subset of twelve strains in high demand was used throughout the study to emphasize various genetic anomalies and patterns (Table 2). The twelve strains were chosen based on popularity (Leafly, 2018; Wikileaf, 2018) and availability.

**Table 1.**
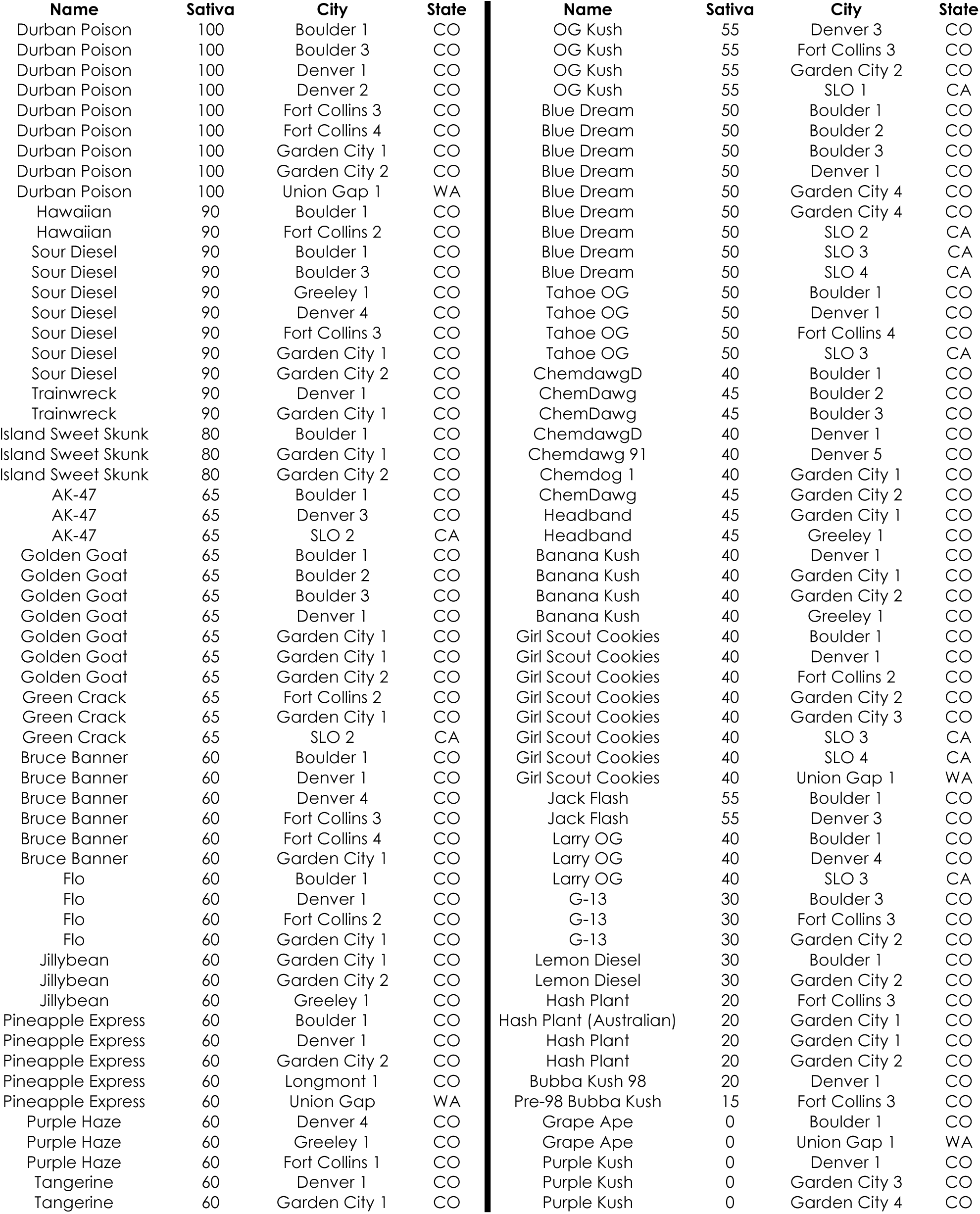
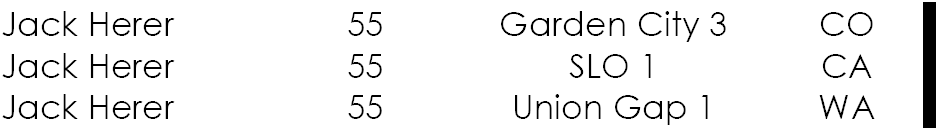
*Cannabis* samples (122) from 30 strains with the reported proportion of Sativa from Wikileaf (Wikileaf, 2018) and the city location and state where each sample was acquired. (SLO: San Luis Obispo).

| Name | Sativa | City | State | Name | Sativa | City | State |
| --- | --- | --- | --- | --- | --- | --- | --- |
| Durban Poison | 100 | Boulder 1 | CO | OG Kush | 55 | Denver 3 | CO |
| Durban Poison | 100 | Boulder 3 | CO | OG Kush | 55 | Fort Collins 3 | CO |
| Durban Poison | 100 | Denver 1 | CO | OG Kush | 55 | Garden City 2 | CO |
| Durban Poison | 100 | Denver 2 | CO | OG Kush | 55 | SLO 1 | CA |
| Durban Poison | 100 | Fort Collins 3 | CO | Blue Dream | 50 | Boulder 1 | CO |
| Durban Poison | 100 | Fort Collins 4 | CO | Blue Dream | 50 | Boulder 2 | CO |
| Durban Poison | 100 | Garden City 1 | CO | Blue Dream | 50 | Boulder 3 | CO |
| Durban Poison | 100 | Garden City 2 | CO | Blue Dream | 50 | Denver 1 | CO |
| Durban Poison | 100 | Union Gap 1 | WA | Blue Dream | 50 | Garden City 4 | CO |
| Hawaiian | 90 | Boulder 1 | CO | Blue Dream | 50 | Garden City 4 | CO |
| Hawaiian | 90 | Fort Collins 2 | CO | Blue Dream | 50 | SLO 2 | CA |
| Sour Diesel | 90 | Boulder 1 | CO | Blue Dream | 50 | SLO 3 | CA |
| Sour Diesel | 90 | Boulder 3 | CO | Blue Dream | 50 | SLO 4 | CA |
| Sour Diesel | 90 | Greeley 1 | CO | Tahoe OG | 50 | Boulder 1 | CO |
| Sour Diesel | 90 | Denver 4 | CO | Tahoe OG | 50 | Denver 1 | CO |
| Sour Diesel | 90 | Fort Collins 3 | CO | Tahoe OG | 50 | Fort Collins 4 | CO |
| Sour Diesel | 90 | Garden City 1 | CO | Tahoe OG | 50 | SLO 3 | CA |
| Sour Diesel | 90 | Garden City 2 | CO | ChemdawgD | 40 | Boulder 1 | CO |
| Trainwreck | 90 | Denver 1 | CO | ChemDawg | 45 | Boulder 2 | CO |
| Trainwreck | 90 | Garden City 1 | CO | ChemDawg | 45 | Boulder 3 | CO |
| Island Sweet Skunk | 80 | Boulder 1 | CO | ChemdawgD | 40 | Denver 1 | CO |
| Island Sweet Skunk | 80 | Garden City 1 | CO | Chemdawg 91 | 40 | Denver 5 | CO |
| Island Sweet Skunk | 80 | Garden City 2 | CO | Chemdog 1 | 40 | Garden City 1 | CO |
| AK-47 | 65 | Boulder 1 | CO | ChemDawg | 45 | Garden City 2 | CO |
| AK-47 | 65 | Denver 3 | CO | Headband | 45 | Garden City 1 | CO |
| AK-47 | 65 | SLO 2 | CA | Headband | 45 | Greeley 1 | CO |
| Golden Goat | 65 | Boulder 1 | CO | Banana Kush | 40 | Denver 1 | CO |
| Golden Goat | 65 | Boulder 2 | CO | Banana Kush | 40 | Garden City 1 | CO |
| Golden Goat | 65 | Boulder 3 | CO | Banana Kush | 40 | Garden City 2 | CO |
| Golden Goat | 65 | Denver 1 | CO | Banana Kush | 40 | Greeley 1 | CO |
| Golden Goat | 65 | Garden City 1 | CO | Girl Scout Cookies | 40 | Boulder 1 | CO |
| Golden Goat | 65 | Garden City 1 | CO | Girl Scout Cookies | 40 | Denver 1 | CO |
| Golden Goat | 65 | Garden City 2 | CO | Girl Scout Cookies | 40 | Fort Collins 2 | CO |
| Green Crack | 65 | Fort Collins 2 | CO | Girl Scout Cookies | 40 | Garden City 2 | CO |
| Green Crack | 65 | Garden City 1 | CO | Girl Scout Cookies | 40 | Garden City 3 | CO |
| Green Crack | 65 | SLO 2 | CA | Girl Scout Cookies | 40 | SLO 3 | CA |
| Bruce Banner | 60 | Boulder 1 | CO | Girl Scout Cookies | 40 | SLO 4 | CA |
| Bruce Banner | 60 | Denver 1 | CO | Girl Scout Cookies | 40 | Union Gap 1 | WA |
| Bruce Banner | 60 | Denver 4 | CO | Jack Flash | 55 | Boulder 1 | CO |
| Bruce Banner | 60 | Fort Collins 3 | CO | Jack Flash | 55 | Denver 3 | CO |
| Bruce Banner | 60 | Fort Collins 4 | CO | Larry OG | 40 | Boulder 1 | CO |
| Bruce Banner | 60 | Garden City 1 | CO | Larry OG | 40 | Denver 4 | CO |
| Flo | 60 | Boulder 1 | CO | Larry OG | 40 | SLO 3 | CA |
| Flo | 60 | Denver 1 | CO | G-13 | 30 | Boulder 3 | CO |
| Flo | 60 | Fort Collins 2 | CO | G-13 | 30 | Fort Collins 3 | CO |
| Flo | 60 | Garden City 1 | CO | G-13 | 30 | Garden City 2 | CO |
| Jillybean | 60 | Garden City 1 | CO | Lemon Diesel | 30 | Boulder 1 | CO |
| Jillybean | 60 | Garden City 2 | CO | Lemon Diesel | 30 | Garden City 2 | CO |
| Jillybean | 60 | Greeley 1 | CO | Hash Plant | 20 | Fort Collins 3 | CO |
| Pineapple Express | 60 | Boulder 1 | CO | Hash Plant (Australian) | 20 | Garden City 1 | CO |
| Pineapple Express | 60 | Denver 1 | CO | Hash Plant | 20 | Garden City 1 | CO |
| Pineapple Express | 60 | Garden City 2 | CO | Hash Plant | 20 | Garden City 2 | CO |
| Pineapple Express | 60 | Longmont 1 | CO | Bubba Kush 98 | 20 | Denver 1 | CO |
| Pineapple Express | 60 | Union Gap | WA | Pre-98 Bubba Kush | 15 | Fort Collins 3 | CO |
| Purple Haze | 60 | Denver 4 | CO | Grape Ape | 0 | Boulder 1 | CO |
| Purple Haze | 60 | Greeley 1 | CO | Grape Ape | 0 | Union Gap 1 | WA |
| Purple Haze | 60 | Fort Collins 1 | CO | Purple Kush | 0 | Denver 1 | CO |
| Tangerine | 60 | Denver 1 | CO | Purple Kush | 0 | Garden City 3 | CO |
| Tangerine | 60 | Garden City 1 | CO | Purple Kush | 0 | Garden City 4 | CO |

|  |  |  |  |
| --- | --- | --- | --- |
| Jack Herer | 55 | Garden City 3 | CO |
| Jack Herer | 55 | SLO 1 | CA |
| Jack Herer | 55 | Union Gap 1 | WA |

**Table 2.** *Cannabis* samples (122) from 30 strains with the reported proportion of Sativa retrieved from Wikileaf (Wikileaf, 2018). Strains arranged by proportion of Sativa, from reported pure Sativa to pure Indica (which has no reported proportion of Sativa) and the proportions of membership for genotype 1 and genotype 2 from the STRUCTURE (Fig. 2) are reported as a percentage according to the proportion of inferred ancestry. Asterisk indicates the twelve popular strains used in further analyses Diamond indicates clone only strains (SeedFinder, 2018)

| Strain | # Samples | Sativa Percentage | Genotype 1 (% average) | Genotype 2 (% average) | Standard Deviation |
| --- | --- | --- | --- | --- | --- |
| Durban Poison* | 9 | 100 | 86 | 14 | 9.9 |
| Hawaiian | 2 | 90 | 61 | 39 | 27.58 |
| Sour Diesel* | 7 | 90 | 14 | 86 | 53.74 |
| Trainwreck | 2 | 90 | 59 | 41 | 21.92 |
| Island Sweet Skunk | 3 | 80 | 93 | 7 | 9.19 |
| AK-47 | 3 | 65 | 55 | 45 | 7.07 |
| Golden Goat** | 7 | 65 | 68 | 32 | 2.12 |
| Green Crack* | 3 | 65 | 60 | 40 | 3.54 |
| Bruce Banner* | 6 | 60 | 19 | 81 | 28.99 |
| Flo* | 4 | 60 | 38 | 62 | 15.56 |
| Jillybean | 3 | 60 | 73 | 27 | 9.19 |
| Pineapple Express* | 5 | 60 | 62 | 38 | 1.41 |
| Purple Haze | 3 | 60 | 77 | 23 | 12.02 |
| Tangerine | 2 | 60 | 53 | 47 | 4.95 |
| Jack Herer | 3 | 55 | 66 | 34 | 7.78 |
| OG Kush** | 4 | 55 | 28 | 72 | 19.09 |
| Blue Dream** | 9 | 50 | 80 | 20 | 21.21 |
| Tahoe OG | 4 | 50 | 26 | 74 | 16.97 |
| Chemdawg* | 7 | 45 | 9 | 91 | 25.46 |
| Headband | 2 | 45 | 57 | 43 | 8.49 |
| Banana Kush* | 4 | 40 | 52 | 48 | 8.49 |
| Girl Scout Cookies** | 8 | 40 | 25 | 75 | 10.61 |
| Jack Flash | 2 | 40 | 96 | 4 | 39.6 |
| Larry OG | 3 | 40 | 7 | 93 | 23.33 |
| G-13 | 3 | 30 | 50 | 50 | 14.14 |
| Lemon Diesel* | 2 | 30 | 85 | 15 | 38.89 |
| Hash Plant | 4 | 20 | 37 | 63 | 12.02 |
| Pre98-Bubba Kush | 2 | 15 | 7 | 93 | 5.66 |
| Grape Ape | 2 | 0 | 55 | 45 | 38.89 |
| Purple Kush** | 4 | 0 | 29 | 71 | 20.51 |

### Microsatellite Development

The *Cannabis* draft genome from ‘Purple Kush’ (GenBank accession AGQN00000000.1) was scanned for microsatellite repeat regions using MSATCOMMANDER-1.0.8-beta (Faircloth, 2008). Primers were developed *de-novo* flanking thirty microsatellites with 3-6 nucleotide repeat units (Table S1). One primer in each pair was tagged with a 5’ universal sequence (M13, CAGT or T7) so that a matching sequence with a fluorochrome tag could be incorporated via PCR (Schwabe *et al.,* 2013). Ten of the thirty primer pairs produced consistent peaks within the predicted size range and were used for the genetic analyses herein.

### PCR and Data Scoring

Microsatellite loci were amplified in 12 µL reactions using 1.0 µL DNA (10-20 ng/ µL), 0.6 µL fluorescent tag (5 µM; FAM, VIC, or PET), 0.6 µL non-tagged primer (5 µM), 0.6 µL tagged primer (0.5 µM), 0.7 µL dNTP mix (2.5mM), 2.4 µL GoTaq Flexi Buffer (Promega, Madison, WI, USA), 0.06 µL GoFlexi taq polymerase (Promega), 0.06 µL BSA (Bovine Serum Albumin 100X), 0.5-6.0 µL MgCl or MgSO_4_, and 0.48-4.98 µL dH_2_O. Amplified products were combined into multiplexes and diluted with water. Hi-Di formamide and LIZ 500 size standard (Applied Biosystems, Foster City, CA, USA) were added before electrophoresis on a 3730 Genetic Analyzer (Applied Biosystems) at Arizona State University. Fragments were sized using GENEIOUS 8.1.8 (Biomatters Ltd).

### Genetic Statistical Analyses

GENALEX ver. 6.4.1 (Peakall & Smouse, 2006; Peakall & Smouse, 2012) was used to calculate deviation from Hardy–Weinberg equilibrium (HWE). Linkage disequilibrium was tested using GENEPOP ver. 4.0.10 (Raymond & Rousset, 1995; Rousset, 2008). The possibility of null alleles was assessed using MICRO-CHECKER (Van Oosterhout*et al.,* 2004). Genotypes were analyzed using the Bayesian cluster analysis program STRUCTURE ver. 2.4.2 (Pritchard *et al.,* 2000). Burn-in and run-lengths of 50,000 generations were used with ten independent replicates for each STRUCTURE analysis. STRUCTURE HARVESTER (Earl, 2012), which implements the Evanno method (Evanno *et al.,* 2005), was used to determine the *K* value that best describes the number of genetic groups for the data set. GENALEX was used to conduct a Principal Coordinate Analysis (PCoA) to examine variation in the dataset. Lynch & Ritland (Lynch & Ritland, 1999) pairwise genetic relatedness (*r*) values were reported for each sample within a strain using GENALEX. Mean pairwise relatedness (*r*) statistics were calculated between all 122 samples resulting in 7381 pairwise *r*-values showing degrees of relatedness. A genetic pairwise relatedness heat map of the data set was generated in Microsoft EXCEL. For all strains the *r-*mean and standard deviation (SD) was calculated averaging among all samples. Obvious outliers were determined by calculating the lowest *r-*mean and iteratively removing those samples to determine the relatedness among the remaining samples in the subset. A graph was generated for the twelve popular strains to show how the *r-*mean value change within a strain when outliers were removed.

## Results

The microsatellite analyses show genetic inconsistencies in *Cannabis* strains acquired from different facilities. The samples used in this study are drug-type strains and are categorized as Sativa, Indica and Hybrid type according to Wikileaf (Wikileaf, 2018). While some popular strains were widely available, some strains were found only at two dispensaries (Table 1 & 2). Since the aim of the research was not to identify specific locations where strain inconsistencies were found, the names for each dispensary are coded to protect the identity of businesses.

There was no evidence of linkage-disequilibrium when all the samples were treated as a single population. All loci deviate significantly from HWE when all samples were treated as a single population, and all but one locus was monomorphic in at least two strains. All but one locus had excess homozygosity and therefore possibly null alleles. Given the inbred nature and extensive hybridization of *Cannabis*, deviations from neutral expectations are not surprising, and the lack of linkage-disequilibrium indicates that the markers are spanning multiple regions of the genome. There was no evidence of null alleles due to scoring errors.

STRUCTURE HARVESTER calculated high support (ΔK=146.56) for two genetic groups, *K=*2 (Fig. 1). STRUCTURE assignment for all samples is shown in Fig. 2 with the strains ordered by the purported proportions of Sativa phenotype (Wikileaf, 2018) and then alphabetically within each strain by city. A clear genetic distinction between Sativa and Indica types would assign 100% Sativa strains (‘Durban Poison’) to one genotype, and assign 100% Indica strains (‘Purple Kush’) to the other genotype (Table 2, Fig. 2). Division of the genotypes into two genetic groups does not support the commonly described Sativa and Indica phenotypes. For the assigned 100% Sativa type strain ‘Durban Poison’, seven of nine samples show greater than 96% assignment to genotype 1 (blue; Fig. 2). For the assigned 100% Indica type ‘Purple Kush’ three of four samples of show greater than 89% assignment to genotype 2 (yellow; Fig. 2). However, samples of ‘Hawaiian’ (90% Sativa) and ‘Grape Ape’ (100% Indica) do not show consistent patterns of predominant assignment to genotype 1 or 2. Interestingly, ‘Durban Poison’ (100% Sativa, n = 9) and ‘Sour Diesel’ (90% Sativa, n = 7) have 86% and 14% average assignment to genotype 1, respectively. Hybrid strains should result in some proportion of shared ancestry, with assignment to both genotype 1 and 2. The strains ‘Blue Dream’ and ‘Tahoe OG’ are reported as 50-50% Sativa-Indica Hybrid strains, but eight of nine samples of ‘Blue Dream’ show > 80% assignment to genotype 1, and three of four samples of ‘Tahoe OG’ show < 7% assignment to genotype 1.

**Fig. 1.**
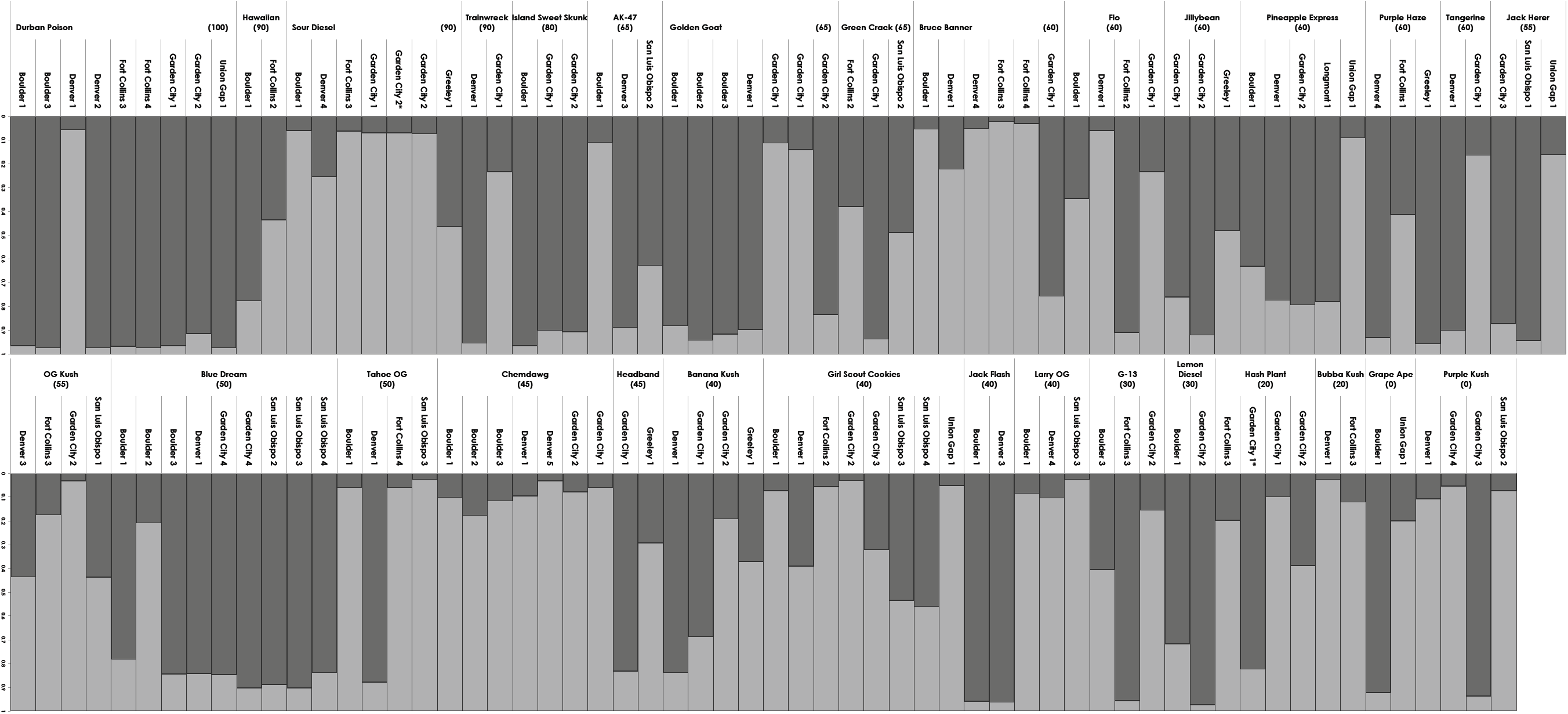
Bar plot graphs generated from STRUCTURE analysis for 122 individuals from 30 strains dividing genotypes into two genetic groups, K=2. Samples were arranged by purported proportions from 100% Sativa to 100% Indica (Wikileaf, 2018) and then alphabetically within each strain by city. Each strain includes reported proportion of Sativa in parentheses (Wikileaf, 2018) and each sample includes the coded location and city from where it was acquired. Each bar indicates proportion of assignment to genotype 1 and genotype 2.

**Fig. 2.**
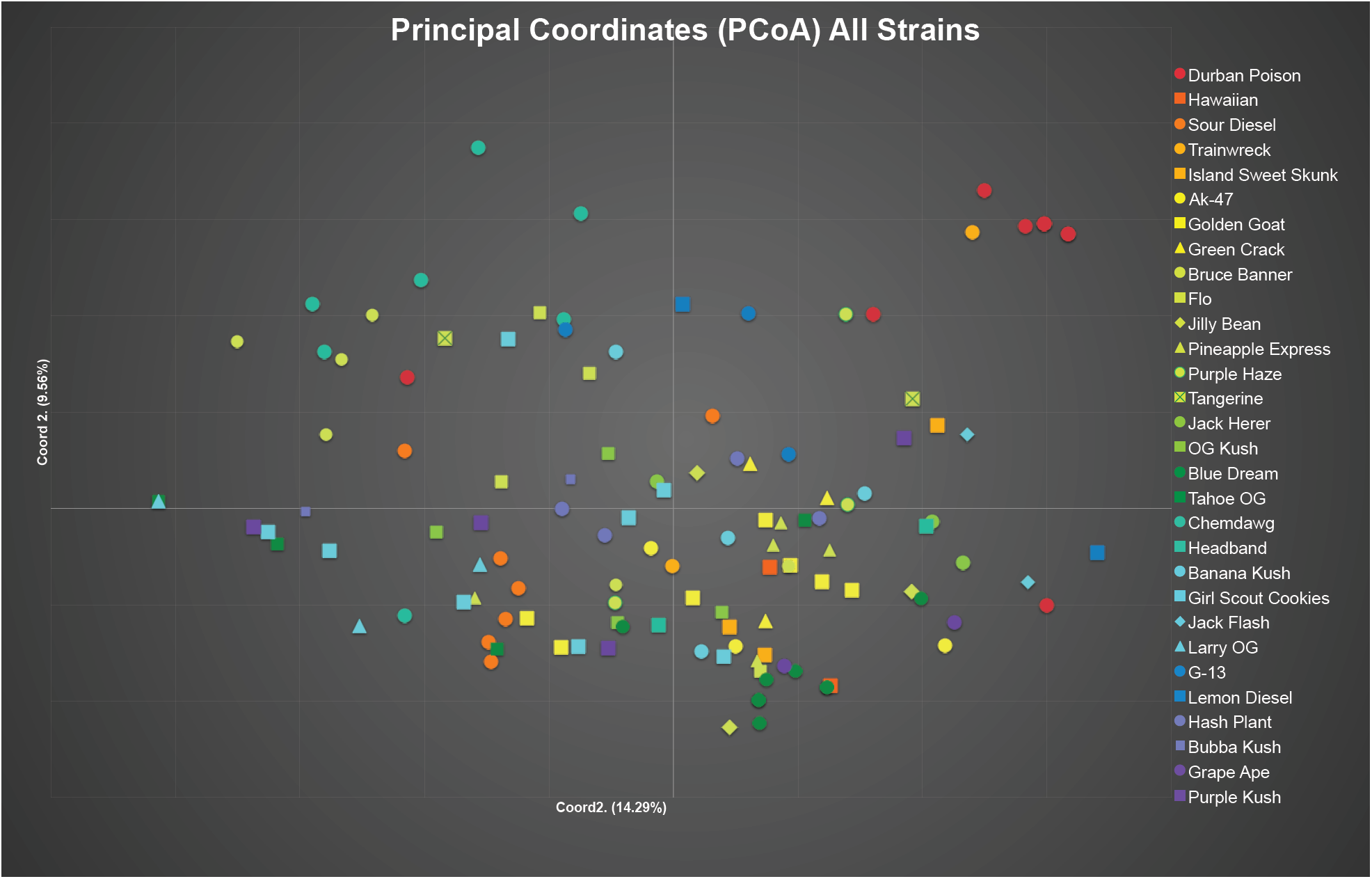
Principal Coordinates Analysis (PCoA) generated in GENALEX. Samples are a color-coded continuum by proportion of Sativa (Table 2) with the strain name given for each sample: Sativa type (red: 100% Sativa proportion, Hybrid type (dark green: 50% Sativa proportion), and Indica type (purple: 0% Sativa proportion). Different symbols are used to indicate different strains within reported phenotype. Coordinate axis 1 explains 14.29% of the variation, coordinate axis 2 explains 9.56% of the variation, and Coordinate axis 3 (not shown) explains 7.07%.

**Fig. 3.**
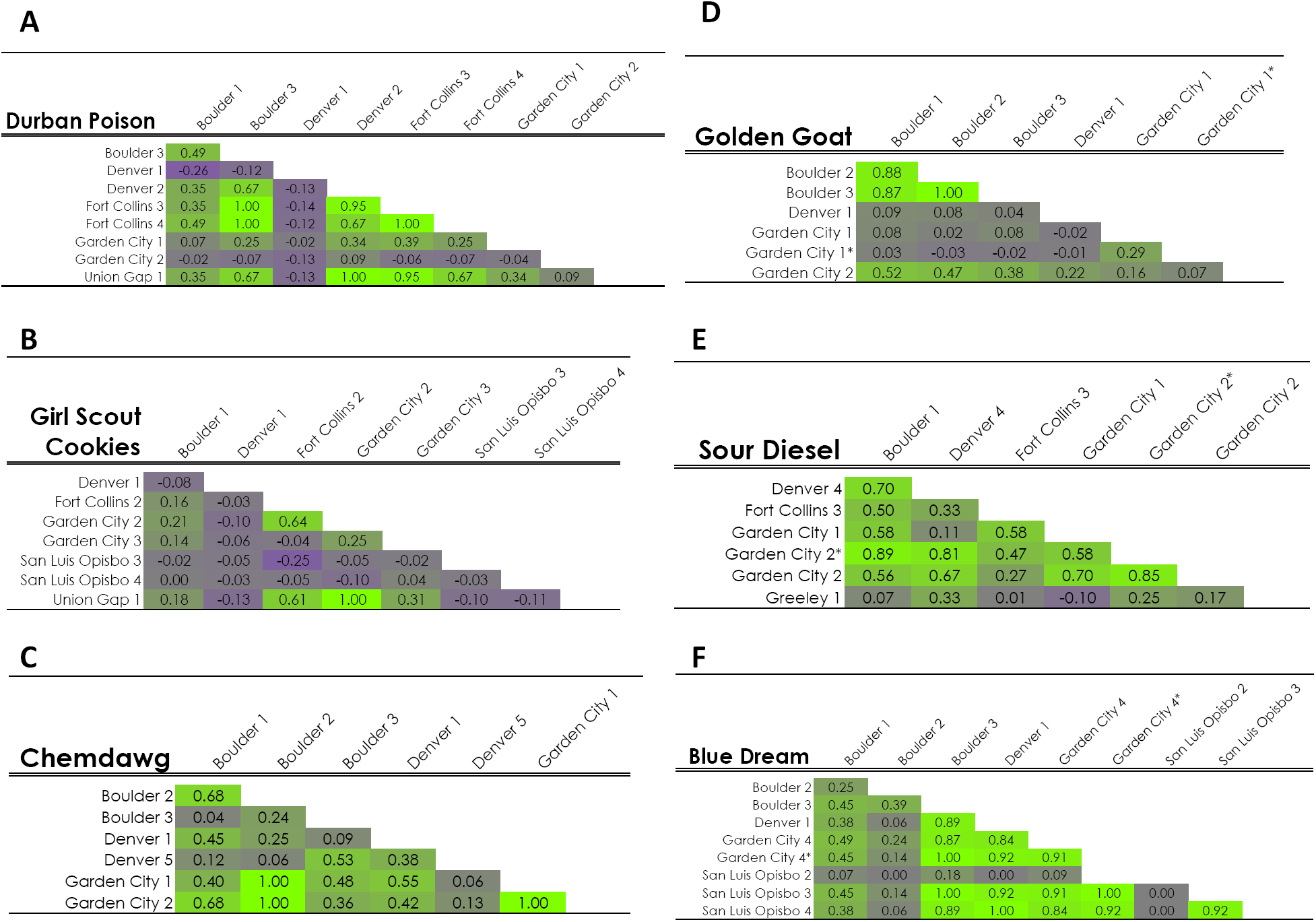
Heat maps of six prominent strains using Lynch & Ritland (1999) pairwise genetic relatedness (*r*) values: purple indicates no genetic relatedness (minimum value −1.09) and green indicates a high degree of relatedness (maximum value 1.0). Sample strain names and location of origin are indicated along the top and down the left side of the chart. Pairwise genetic relatedness (*r*) values are given in each cell and cell color reflects the degree to which two individuals are related.

Principal Coordinate Analyses (PCoA) were conducted using GENALEX for (1) all samples (Fig. 2) and (2) twelve popular strains (Fig. S2). The samples in the PCoA of all 30 strains are organized from 100% Sativa types (red), through all levels of Hybrid types, to 100% Indica types (purple; Fig. 4). Strain types with the same reported proportions are the same color but have different symbols. The PCoA of all strains represents 14.90% of the variation in the data on coordinate axis 1, 9.56% on axis 2, and 7.07% on axis 3 (not shown). The second PCoA of twelve popular strains specifically examines the genetic relationship within strains that are in high demand (Fig. S2). The results from this analysis found that 15.30% of the variation in the data is explained by coordinate axis 1, 12.98% on axis 2, and 7.96% on axis 3 (not shown).

**Fig. 4.**
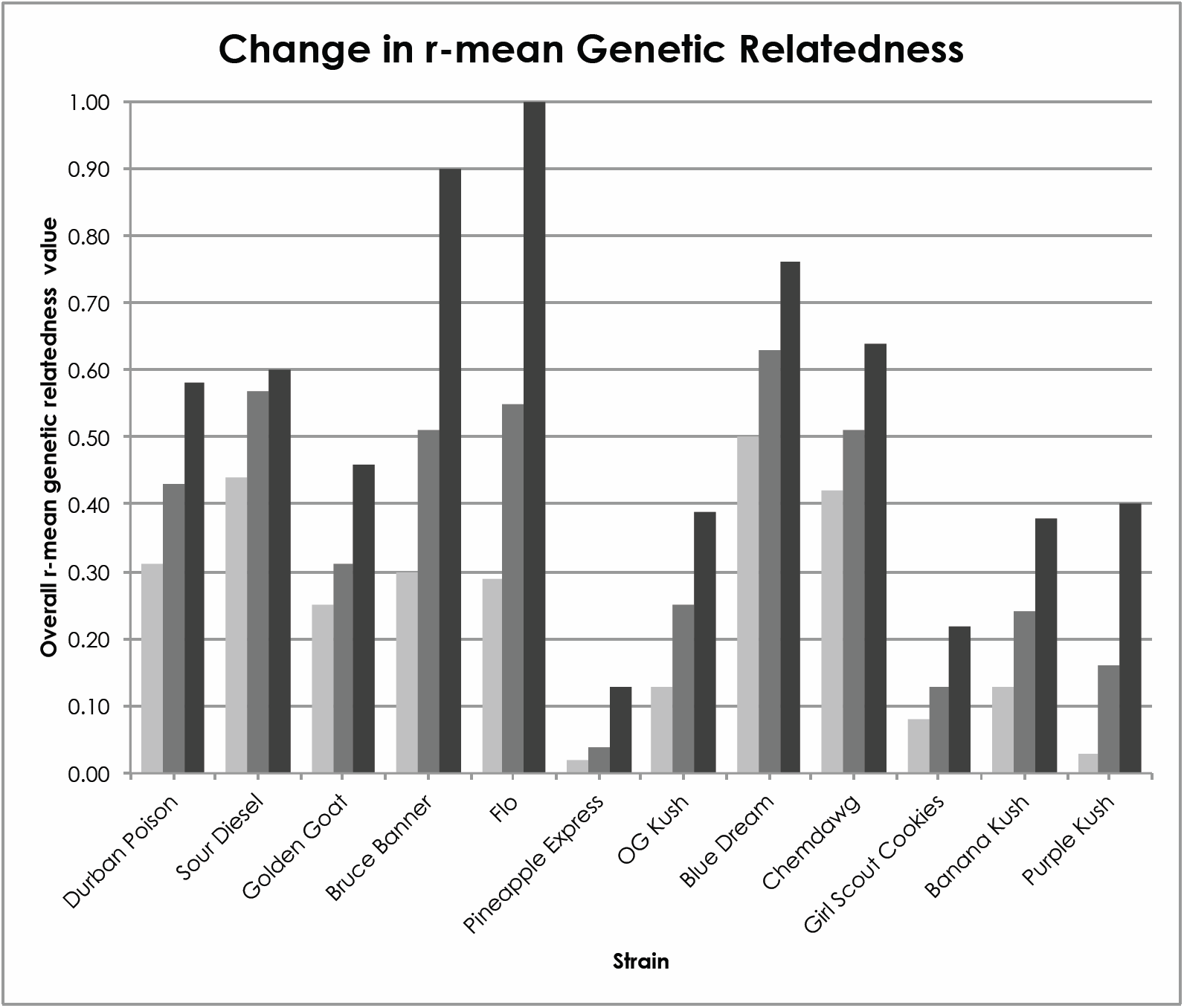
This graph indicates the mean pairwise genetic relatedness (*r*) initially (light gray) and after the removal of one (medium gray) or two (dark gray) outlying samples in 12 prominent strains.

Lynch & Ritland (Lynch & Ritland, 1999) pairwise genetic relatedness (*r*) between all 122 samples was calculated in GENALEX. The resulting 7380 pairwise *r*-values were converted to a heat map using purple to indicate the lowest pairwise relatedness value (−1.09) and green to indicate the highest pairwise relatedness value (1.00; Fig. S3. Comparisons are detailed for six popular strains (Fig. 3) to illustrate the relationship of samples from different sources and the impact of outliers. Values of close to 1.00 indicate a high degree of relatedness (Lynch & Ritland, 1999), which could be indicative of clones or seeds from the same mother (Green, 2005; SeedFinder, 2017). First order relatives (full siblings or mother-daughter) share 50% genetic identity (*r-*value = 0.50), second order relatives (half siblings or cousins) share 25% genetic identity (*r-*value = 0.25), and unrelated individuals are expected to have an *r-*value of 0.00 or lower. Negative values arise when individuals are less related than expected under normal panmictic conditions (Moura *et al.,* 2013; Norman *et al.,* 2017). Values ranged from −1.09 (between ‘Purple Haze’ Greeley 1 and ‘Girl Scout Cookies’ Union Gap 1) indicating low levels of relatedness, to 1.00 (e.g., between ‘Durban Poison’ samples from Boulder 3 and Fort Collins 3).

Individual pairwise *r-*values were averaged within strains to calculate the overall *r-*mean as a measure of genetic similarity within strains. The overall *r-*means within strains ranged from −0.22 (‘Tangerine’) to 0.68 (‘Island Sweet Skunk’) (Table 3). Standard deviations ranged from 0.04 (‘Jack Herer) to 0.51 (‘Bruce Banner’). The strains with higher standard deviation values indicate a wide range of genetic relatedness within a strain, while low values indicate that samples within a strain share similar levels of genetic relatedness. In order to determine how outliers impact the overall relatedness in a strain, the farthest outlier (lowest pairwise *r-*mean value) was removed and the overall *r-*means and SD values within strains were recalculated (Table 3). In all strains, the overall *r-*means increased when outliers were removed. In strains with more than three samples, a second outlier was removed and the overall *r-*means and SD values were recalculated. Overall *r-*means were used to determine degree of relatedness as clonal (or from stable seed; overall *r-*means > 0.9), first or higher order relatives (overall *r-*means 0.46 – 0.89), second order relatives (overall *r-*means 0.26 - 0.45), low levels of relatedness (overall *r-* means 0.00 - 0.25), and not related (overall *r-*means <0.00). Initial overall *r-*means indicate only three strains are first or higher order relatives (Table 3). Removing outliers revealed samples within ten of the remaining 22 strains are first or higher order relatives. After outliers were removed, 15 of the 30 strains are comprised of first or higher order relatives, indicating outliers are often responsible for variability within strains. Removing outliers revealed samples within seven of the twelve popular strains are of first or higher order relatives (Table 3, Fig. 4). Three strains are comprised of second order relatives with overall *r-*means ranging from 0.22 - 0.25. Two strains show low levels of relatedness with overall *r-*means ranging from 0.13 - 0.16 even after outliers are removed (Table 3). The impact of outliers can be clearly seen in the heat map for ‘Durban Poison’ which shows the relatedness for 36 comparisons (Fig. 3A), six of which are nearly identical (*r*-value 0.90 - 1.0), six of which are first order siblings (*r*-value 0.46 - 0.89), six of which are second order relatives (*r*-value 0.26 - 0.45), five of which have low levels of relatedness (*r*-value 0.00 - 0.25), and 13 which are not related (*r*-value <0.00). However, removal of two outliers, Denver 1 and Garden City 2, reduces the number of comparisons ranked as not related from 13 to zero, and low level of relatedness from five to one.

**Table 3.** Lynch & Ritland (1999) pairwise relatedness comparisons of overall *r*-means (Mean) and standard deviations (SD) for samples of 30 strains including *r*-mean and SD after the first and second (where possible) outliers were removed. Outliers were samples with the lowest *r*-mean. The twelve popular strains are indicated with an asterisk. Diamonds indicate clone-only strains (SeedFinder, 2018)

| Strain | # Samples | Measure | All samples | Outlier 1 removed | Outlier 2 removed |
| --- | --- | --- | --- | --- | --- |
| Durban Poison* | 9 | Mean | 0.31 | 0.43 | 0.58 |
|  |  | SD | 0.40 | 0.37 | 0.30 |
| Hawaiian | 2 | Mean | -0.115 | - | - |
|  |  | SD |  |  |  |
| Sour Diesel* | 7 | Mean | 0.44 | 0.57 | 0.60 |
|  |  | SD | 0.29 | 0.22 | 0.18 |
| Trainwreck | 2 | Mean | -0.001 | - | - |
|  |  | SD |  |  |  |
| Island Sweet Skunk | 3 | Mean | 0.682 | 1.000 | - |
|  |  | SD |  |  |  |
| AK-47 | 3 | Mean | 0.158 | 0.446 | - |
|  |  | SD |  |  |  |
| Golden Goat** | 7 | Mean | 0.25 | 0.31 | 0.46 |
|  |  | SD | 0.32 | 0.36 | 0.36 |
| Green Crack* | 3 | Mean | 0.375 | 0.885 | - |
|  |  | SD |  |  |  |
| Bruce Banner* | 6 | Mean | 0.30 | 0.51 | 0.90 |
|  |  | SD | 0.51 | 0.50 | 0.05 |
| Flo* | 4 | Mean | 0.29 | 0.55 | - |
|  |  | SD | 0.38 | 0.39 | - |
| Jillybean | 3 | Mean | -0.033 | 0.039 | - |
|  |  | SD |  |  |  |
| Pineapple Express* | 5 | Mean | 0.02 | 0.04 | 0.13 |
|  |  | SD | 0.16 | 0.17 | 0.19 |
| Purple Haze | 3 | Mean | 0.041 | 0.263 | - |
|  |  | SD |  |  |  |
| Tangerine | 2 | Mean | -0.219 | - | - |
|  |  | SD |  |  |  |
| Jack Herer | 3 | Mean | 0.102 | 0.127 | - |
|  |  | SD |  |  |  |
| OG Kush** | 4 | Mean | 0.13 | 0.25 | - |
|  |  | SD | 0.19 | 0.22 | - |
| Blue Dream** | 9 | Mean | 0.50 | 0.63 | 0.76 |
|  |  | SD | 0.39 | 0.34 | 0.24 |
| Tahoe OG | 4 | Mean | 0.210 | 0.406 | 0.539 |
|  |  | SD |  |  |  |
| Chemdawg* | 7 | Mean | 0.42 | 0.51 | 0.64 |
|  |  | SD | 0.31 | 0.31 | 0.28 |
| Headband | 2 | Mean | 0.107 | - | - |
|  |  | SD |  |  |  |
| Banana Kush* | 4 | Mean | 0.13 | 0.24 | - |
|  |  | SD | 0.20 | 0.13 | - |
| Girl Scout Cookies** | 8 | Mean | 0.08 | 0.13 | 0.22 |
|  |  | SD | 0.27 | 0.30 | 0.32 |
| Jack Flash | 2 | Mean | 0.621 | - | - |
|  |  | SD |  |  |  |
| Larry OG | 3 | Mean | 0.316 | 0.671 | - |
|  |  | SD |  |  |  |
| G-13 | 3 | Mean | 0.286 | 0.562 | - |
|  |  | SD |  |  |  |
| Lemon Diesel* | 2 | Mean | 0.102 | - | - |
|  |  | SD |  |  |  |
| Hash Plant | 4 | Mean | 0.250 | 0.250 | 0.427 |
|  |  | SD |  |  |  |
| Pre98-Bubba Kush | 2 | Mean | -0.024 | - | - |
|  |  | SD |  |  |  |
| Grape Ape | 2 | Mean | -0.050 | - | - |
|  |  | SD |  |  |  |
| Purple Kush** | 4 | Mean | 0.03 | 0.16 | - |
|  |  | SD | 0.21 | 0.22 | - |

## Discussion

The legal status and social attitudes toward *Cannabis* are changing worldwide, with more than half the states in the U.S. having sanctioned medical *Cannabis* use (ProCon, 2016a). *Cannabis* types and strains are becoming an ever-increasing topic of discussion, so it is important that scientists and the public can discuss *Cannabis* in a similar manner. Currently, not only are Sativa and Indica types disputed, but also experts are at odds about nomenclature for *Cannabis* (Clarke & Merlin, 2015; Small, 2015b). We investigated the possibility of a genetic distinction in commonly described Sativa and Indica strains. Previous genetic research found genetic variability among seeds from the same strain supplied from a single source, indicating genotypes within strains are variable (Sohler *et al.,* 2017). However, it was unclear if the seeds in the study were produced from multiple parent plants, which could have introduced a source for genetic variation. The focus of this study is that genetic profiles from strains with the same identifying name should have identical, or at least, highly similar genotypes no matter the source of origin. It is important that strain names reflect consistent genetic identity, especially for those who rely on *Cannabis* to alleviate specific medical symptoms. An important element for this study is that samples were acquired from multiple locations to maximize the potential for variation among samples. The multiple genetic analyses used here address important questions and bring scientific evidence to support claims that inconsistent products are being distributed. Genotype analysis can be used to ensure higher levels of consistency within strains. Maintenance of the genetic integrity of strains is possible only following evaluation of genetic consistency, and continuing to overlooking this aspect will to promote variability and phenotypic variation. Addressing strain variability at the molecular level is of the utmost importance while the industry is still relatively new.

Genetic analyses have consistently found genetic distinction between hemp and marijuana, but no clear distinction has been shown between the common description of Sativa and Indica types (de Meijer *et al.,* 1996; Small, 1997; Lynch *et al.,* 2016; Sawler *et al.,* 2015; Vergara *et al.,* 2016; Dufresnes *et al.,* 2017; Soler *et al.,* 2017). We found high support for two genetic groups in the data (Fig. 1) but no discernable distinction or pattern between the described Sativa and Indica strains. The color-coding of strains in the PCoA for all 122 samples allows for visualization of clustering among similar phenotypes by color Sativa (red/orange), Indica (blue/purple) and Hybrid (green) type strains (Fig. 2). However, there is no evidence of clustering in the three commonly described types. If genetic differentiation of the commonly perceived Sativa and Indica types previously existed, it is no longer detectable in the neutral genetic markers used here. Extensive hybridization and selection has presumably created a homogenizing effect and erased evidence of potentially divergent historical genotypes.

Wikileaf maintains that the proportions of Sativa and Indica reported for strains are largely based on genetics and lineage (Dan Nelson, Wikileaf, personal communication). This has seemingly become convoluted over time (Russo, 2007; Small, 2015a; Clarke & Merlin, 2013; Small, 2017). Our results show that commonly reported levels of Sativa, Indica and Hybrid type strains are often not reflected in the average genotype. For example, two sought-after Sativa strains, ‘Durban Poison’ and ‘Sour Diesel’, were found to have contradicting genetic assignments (Fig. 1, Table 2). ‘Durban Poison’, described as 100% Sativa, has an 86% average assignment to genotype 1, while ‘Sour Diesel’, described as 90% Sativa, has a 14% average assignment to genotype 1. This analysis indicates strains with similar reported proportions of Sativa or Indica may have differing genetic assignments. Further illustrating this point is that ‘Bruce Banner’, ‘Flo’, ‘Jillybean’, ‘Pineapple Express’, ‘Purple Haze’, and ‘Tangerine’ are all reported to be 60/40 Hybrid type strains, but clearly have differing levels of admixture both within and among these reportedly similar strains (Table 2, Fig. 1). From these results, we can conclude that reported ratios or differences between Sativa and Indica phenotypes are not discernable using these genetic markers. Given the lack of genetic distinction between Indica and Sativa types, it is not surprising that reported ancestry proportions are also not supported.

To accurately address reported variation within strains, samples were purchased from various locations, as a customer, with no information of strains other than publically available online information. Evidence for genetic inconsistencies is apparent within many strains and supported by multiple genetic analyses. In our analyses of 30 strains, only 4 strains had consistent STRUCTURE genotype assignment and admixture among all samples: ‘Chemdawg’ (n=7), ‘Island Sweet Skunk’ (n=3), ‘Larry OG’ (n=3) and ‘Jack Flash’ (n = 2; Fig. 2). However, it is clear that many strains contained one or more obvious genetic outliers (e.g. Durban Poison – Denver 1; Fig 1, 3A). With the removal of one obvious outlier, the remaining samples of eleven strains were classified as first order relatives based on pairwise genetic relatedness r-values (overall *r-*mean >0.45; Table 3, Fig. 4). The removal of a second outlier resulted in 15 of the 30 strains having an overall *r-*mean >0.45 (Table 3, Fig. 4). Together, these results indicate that half of the strains used in this analysis showed relatively stable genetic identity among most samples within a strain. Six of the strains with inconsistent patterns had only two samples, both of which were different (e.g., ‘Trainwreck’ and ‘Headband’). The remaining nine strains in the analysis had more than one obvious outlier (e.g., ‘Sour Diesel’) or had no consistent genetic pattern among the samples within the strain (e.g., ‘Girl Scout Cookies’; Table 3, Fig. 1, Fig. 2, Fig. S2). It is noteworthy that many of the strains used here fell into a range of genetic relatedness indicative of first order siblings (*r*-value 0.46 - 0.89) when samples with high genetic divergence were isolated and removed from the data set (Table 4; Figs. 3, 4).

Relationships within the twelve popular strains were analyzed separately to determine if (1) strains with more samples show a higher degree of clustering, and (2) strains in higher demand have a higher degree of genetic relatedness. The analysis of genetic variation for the subset of twelve popular strains shows some clustering within strains (Fig. S2), but clustering is not seen for all strains, and outliers are apparent. This analysis represents more of the variation in the data compared to the PCoA for all 30 strains and shows clustering of some strains, such as ‘Durban Poison’, ‘Golden Goat’ and ‘Blue Dream’. However, all clusters have at least one sample that is removed from the other samples in the group. From this we argue that samples representing the popular strains may be slightly more likely to have a higher degree of genetic relatedness, but more sampling would be required to determine this with confidence.

A pairwise genetic heat map based on Lynch & Ritland (Lynch & Ritland, 1999) pairwise genetic relatedness (*r-*values) was generated to visualize genetic relatedness throughout the data set (Fig. S3). Values of 1.00 (or close to) are assumed to be clones or plants from self-fertilized seed. Six examples of within-strain pairwise comparison heat maps were examined to illustrate common patterns (Fig.7). The heat map shows that many strains contain samples that are first order relatives or higher (*r*-value > 0.49). For example ‘Sour Diesel’ (Fig. 3??) has 12 comparisons of first order or above, and six have low/no relationship. There are also values that could be indicative of clones or plants from a stable seed source such as ‘Blue Dream’ (Fig. 3???), which has 10 nearly identical comparisons (*r*-value 0.90-1.00), and no comparisons in ‘Blue Dream’ have negative values. While ‘Blue Dream’ has an initial overall *r*-mean indicating first order relatedness within the samples (Table 3, Fig. 4), it still contains more variation than would be expected from a clone only strain (SeedFinder, 2017). Other clone-only strains (SeedFinder, 2017), e.g. ‘Girl Scout Cookies’ (Table 3, Fig. 3??) and ‘Golden Goat’ (Table 3, Fig. 3??), have a high degree of genetic variation resulting in low overall relatedness values. Outliers were calculated and removed iteratively to demonstrate how they affected the overall *r*-mean within the twelve popular strains (Table 3, Fig. 4). In all cases, removing outliers increased the mean *r*-value, as illustrated by ‘Bruce Banner’, which increased substantially, from 0.3 to 0.9 when samples with two outlying genotypes removed. The outliers are evidence of inconsistencies within strains and when removed, genetic relatedness greatly improves. There are unexpected areas in the heat map that indicate high degrees of relatedness between different strains (Fig. S3). For example, comparisons between ‘Golden Goat’ and ‘Island Sweet Skunk’ (overall *r*-mean 0.37) are higher than within samples of ‘Sour Diesel’. Interestingly, ‘Golden Goat’ is reported to be a hybrid descendant of ‘Island Sweet Skunk’ (Leafly, 2018), which explains the high genetic relatedness between these strains. However, most of the between strain overall *r*-mean are negative (e.g., ‘Golden Goat’ to ‘Durban Poison’ −0.03 and ‘Chemdawg’ to ‘Durban Poison’ −0.22; Fig. S3), indicative of limited recent genetic relationship.

While collecting samples from various dispensaries, it was noted that strains of ‘Chemdawg’ had various different spellings of the strain name, as well as numbers and/or letters attached to the name. Without knowledge of the history of ‘Chemdawg’, the assumption was that these were local variations. These were acquired to include in the study to determine if and how these variants were related. Upon investigation of possible origins of ‘Chemdawg’, an interesting history was uncovered, especially in light of the results (Backes & Weil, 2014). Legend has it that someone named “Chemdog” (a person) grew the variations (‘Chemdawg 91’, ‘Chemdawg D’, ‘Chemdawg 4’, ‘Chemdog 1’) from seeds he found in an ounce he purchased at a Grateful Dead concert. This illustrates how *Cannabis* strains may have come to market in a non-traditional manner. The history of ‘Chemdawg’ is currently unverifiable, but the analysis supports that these variations could be from seeds of the same plant. Genetic analyses can add scientific support to the stories behind vintage strains and possibly help clarify the history of specific strains.

Possible facilitation of inconsistencies may come from both suppliers and growers of *Cannabis* clones and stable seed, because currently they can only assume the strains they possess are true to name. There is a chain of events from seed to sale that relies heavily on the supplier, grower, and dispensary to provide the correct product, but there is currently no reliable way to verify *Cannabis* strains. The possibility exists for errors in plant labeling, misplacement, misspelling, and/or relabeling along the entire chain of production. Although the expectation is that plants are labeled carefully and not re-labeled with a more desirable name for a quick sale, these misgivings must be considered. Identification by genetic markers has largely eliminated these types of mistakes in other widely cultivated crops such as grapes, olives and apples. Modern genetic applications can accurately identify varieties and can clarify ambiguity in closely related and hybrid species, [e.g., Rongwen *et al.,* 1995; Guilford *et al.,* 1997; Belaj *et al*. 2004; Muzzalupo *et al.,* 2009; Štajner *et al.,* 2011).

Matching genotypes within the same strains were expected, but highly similar genotypes between samples of different strains could be the result of mislabeling or misidentification, especially when acquired from the same source. The pairwise genetic relatedness *r-*values were examined for incidence of possible mislabeling or re-labeling. There were instances in which different strains had *r*-values = 1.0 (Fig. S3), indicating clonal genetic relationships. Two samples with matching genotypes were obtained from the same location (‘Larry OG’ and ‘Tahoe OG’ from San Luis Obispo 3). This could be evidence for mislabeling or misidentification because these two samples have similar names. It is unlikely that these samples from reportedly different strains have identical genotypes, and more likely that these samples were mislabeled at some point. Misspelling may also be a source of error, especially when facilities are handwriting labels. An example of possible misspelling may have occurred in the sample labeled ‘Chemdog 1’ from Garden City 1. ‘Chemdawg 1’, a described strain, could have easily been misspelled, but it is unclear whether this instance is evidence for mislabeling or renaming a local variant. Inadvertent mistakes may carry through to scientific investigation where strains are spelled or labeled incorrectly. For example, Vergara et al. (2016) reports genome assemblies for ‘Chemdog’ and ‘Chemdog 91’ as they are reported in GenBank (GCA_001509995.1), but neither of these labels are recognized strain names. It is likely that these are ‘Chemdawg’ and ‘Chemdawg 91’ (Leafly, 2018; Wikileaf, 2018) although it is possible these strains are unreported variants. Another example that may lead to confusion is how information is reported in public databases. For example, data is available for the reported monoisolate of ‘Pineapple Banana Bubba Kush’ in GenBank (SAMN06546749), and while ‘Pineapple Kush’, ‘Banana Kush’ and ‘Bubba Kush’ are known strains (Leafly, 2018; Wikileaf, 2018), the only record of ‘Pineapple Banana Bubba Kush’ is in Genbank. This study has highlighted several possible sources of error and how genotyping can serve to uncover sources of variation. Although this study was unable to confirm sources of error, it is important that producers, growers and consumers are aware that there are errors and they should be documented and corrected whenever possible.

## Conclusion

Over the last decade, the legal status of *Cannabis* has shifted and is now legal for medical use, and some recreational adult use, in the majority of the United States as well as several other countries that have legalized or decriminalized *Cannabis*. The recent legal changes have led to an unprecedented increase in the number of strains available to consumers. There are currently no baseline genotypes for any strains, but steps should be taken to ensure products marketed as a particular strain are genetically congruent. Although the sampling in this study was not exhaustive, the results are clear: strain inconsistency is evident and is not limited to a single source, but rather exists among dispensaries across cities in multiple states. Various suggestions for naming the genetic variants do not seem to align with the current widespread definitions of Sativa, Indica, Hybrid, and Hemp (Hillig, 2005; Clarke & Merlin, 2013). As our *Cannabis* knowledge base grows, so does the communication gap between scientific researchers and the public. Currently, there is no way for *Cannabis* suppliers, growers or consumers to definitively verify strains. Exclusion from protection, due to the Federal status of *Cannabis* as a Schedule I drug, has created avenues for error and inconsistencies. Presumably, the genetic inconsistencies will often manifest as differences in overall effects (Backes, 2014). Differences in characteristics within a named strain may be surprising for a recreational user, but differences may be more serious for a medical patient who relies on a particular strain for alleviation of specific symptoms.

This study shows that in neutral genetic markers, there is no consistent genetic differentiation between the widely held perceptions of Sativa and Indica *Cannabis* types. Moreover, the genetic analyses do not support the reported proportions of Sativa and Indica within each strain, which is expected given the lack of genetic distinction between Sativa and Indica. Instances were found where samples within strains are not genetically similar, which is unexpected given the manner in which *Cannabis* plants are propagated. Although it is impossible to determine the source of these inconsistencies as they can arise at multiple points throughout the chain of events from seed to sale, we theorize misidentification, mislabeling, misplacement, misspelling, and/or relabeling are all possible. Especially where names are similar, there is the possibility for mislabeling, as was shown here. In many cases genetic inconsistencies within strains were limited to one or two samples. We feel that there is a reasonable amount of genetic similarity within many strains, but currently there is no way to verify the “true” genotype of any strain. Although the sampling here includes merely a fragment of the available *Cannabis* strains, our results give scientific merit to claims that strains can be unpredictable.

## Supplementary Data

Table S1: Primer information used in this research.

Fig. S1: STRUCTURE HARVESTER graph indicating K=2 is highly supported.

Fig. S2: Principal Coordinates Analysis (PCoA) for twelve popular strains.

Fig. S3: Pairwise genetic relatedness (*r*) heat table with values for 122 samples.

## Acknowledgements

We thank Gerald Bresowar and Nolan Kane for comments on an earlier draft of this manuscript. The University of Northern Colorado School of Biological Sciences supported this research, and we are grateful to the Graduate Student Association and the Gerald Schmidt Memorial Biology Scholarship for providing partial funding to carry out this research.

## List of abbreviations

US: United States
HIV: human immunodeficiency virus
AIDS: acquired immune deficiency syndrome
PTSD: post-traumatic stress disorder
THC: Δ^*9*^-tetrahydrocannabinol
USDA: United States Department of Agriculture
PVPA: The Plant Variety Protection Act
PVPO: Plant Variety Protection Office
SLO: San Luis Obispo
DNA: deoxyribonucleic acid
CTAB: Acetyl trimethylammonium bromide
PCR: Polymerase chain reaction
HWE: Hardy–Weinberg equilibrium
PCoA: Principle Coordinates Analysis
SD: standard Deviation
IA: identical alleles

